# Note on a Small Chlamyphoridae (Xenarthra, Cingulata) from the Amazon, Brazil

**DOI:** 10.64898/2026.10.26.762017

**Authors:** Artur Chahud

## Abstract

The Awá-Guajá are one of the last hunter-gatherer peoples in the world and live in Indigenous reserves in northwestern Maranhão, Brazil. The Guajá Ethnographic Collection consists of animal remains found during excavations of the community’s refuse areas, conducted between 1987 and 1990. This study presents a taxonomic reassessment of a small Cingulata specimen from this collection. The specimen, previously identified as *Euphractus sexcinctus*, consists of a complete right femur whose taxonomic identity was reevaluated through comparative morphological and metric analyses using specimens from scientific collections and data from the specialized literature. The results revealed that the femur is smaller than those observed in individuals of *E. sexcinctus* and exhibits a combination of anatomical features resembling those found in the genera *Chaetophractus, Cabassous*, and *Euphractus*. Despite these similarities, the observed morphological differences do not allow a reliable identification at the genus level. Therefore, the specimen was provisionally assigned to the family Chlamyphoridae, and its identification was retained at the family level.

## INTRODUCTION

Armadillos belong to the order Cingulata, one of the main branches of the superorder Xenarthra, a group of placental mammals whose evolutionary history unfolded predominantly in South America.

The diversification of armadillos occurred in the context of South America’s relative geographic isolation, long-term climatic changes, the diversification of terrestrial environments, and, subsequently, faunal exchange with North America. The order Cingulata exhibits considerable morphological diversity, ranging from small, burrowing armadillos to large pampatheres and glyptodonts. The dermal carapace, formed by osteoderms (ossifications that develop in the dermis), is one of the most distinctive characteristics of the group (Gaudin & Croft, 2015; Delsuc et al., 2016; Casali et al., 2026).

The family Chlamyphoridae was established in modern classifications based on molecular evidence demonstrating that the armadillos traditionally grouped within Dasypodidae do not constitute a monophyletic group. Chlamyphoridae includes the subfamilies Euphractinae, Tolypeutinae, and Chlamyphorinae, as well as the glyptodont lineage, according to phylogenetic classifications that recognize the evolutionary proximity of this group to modern armadillos (Delsuc et al., 2016).

The family comprises eight genera: *Euphractus* Wagler, 1830; *Zaedyus* Ameghino, 1889; *Chaetophractus* Fitzinger, 1871; *Chlamyphorus* Harlan, 1825; *Calyptophractus* Fitzinger, 1871; *Priodontes* F. Cuvier, 1825; *Tolypeutes* Illiger, 1811; and *Cabassous* McMurtrie, 1831 (Abba et al., 2015; Delsuc et al., 2016; Gibb et al., 2016; Casali et al., 2026).

The Awá-Guajá are an Indigenous hunter-gatherer people who inhabit the eastern Amazon, in the northwestern part of the state of Maranhão, within the basins of the Pindaré and Gurupi rivers (Garcia, 2010). They refer to themselves as “Awá,” a term meaning “person,” “people,” or “man,” and are believed to have originated in the lower Tocantins River region, in the state of Pará (Balée, 1994). Since the late 1980s, the Awá-Guajá have been distributed across three Indigenous Territories (ITs), within which the National Indian Foundation (FUNAI) has registered the village communities: Alto Turiaçu (530,525 ha), Awá (118,000 ha), and Caru (172,667 ha) (Oliveira et al., 2011a; Prado et al., 2012).

The Guajá Ethnographic Collection (also referred to as the Guajá Collection) consists of faunal remains from the Caru Indigenous Territory in Maranhão, accumulated near an Indigenous encampment between 1987 and 1990 (Queiroz & Kipnis, 1997). This collection provides information on local hunting and gathering practices before the introduction and adoption of agriculture by the community, a process that partially altered these practices (Queiroz & Kipnis, 1997; Prado et al., 2012). In addition, the material provides evidence for reconstructing the composition of the local fauna and documenting its occurrence in the late 1980s (Chahud, 2019, 2020a, 2020b, 2021b; Chahud & Okumura, 2020; Ferreira Figueiredo et al., 2022; Praita et al., 2023a, 2023b).

Among the studies on Xenarthra based on material from the Awá-Guajá collection, Chahud (2022) stands out for identifying two sloth species, one anteater species, specimens of Dasypodidae, and one specimen of *Euphractus sexcinctus*. However, subsequent observations revealed that the latter specimen is considerably smaller than adult individuals and even some subadults of *E. sexcinctus*, indicating the need to reassess its taxonomic identification.

Therefore, the present study aims to provide a detailed redescription of this specimen, whose size and morphology differ from those of other cingulates recorded in the region.

## MATERIAL AND METHODS

The material analyzed consists of a femur from an adult individual, cataloged under number E27S18/09, recovered from food remains associated with the Awá-Guajá community in the Caru Indigenous Territory, located on the banks of the Pindaré River in the state of Maranhão (Figure 1). The specimen was collected during the dry season, between July and August 1990, in areas adjacent to dwellings (Queiroz & Kipnis, 1997).

**Figure 1.**
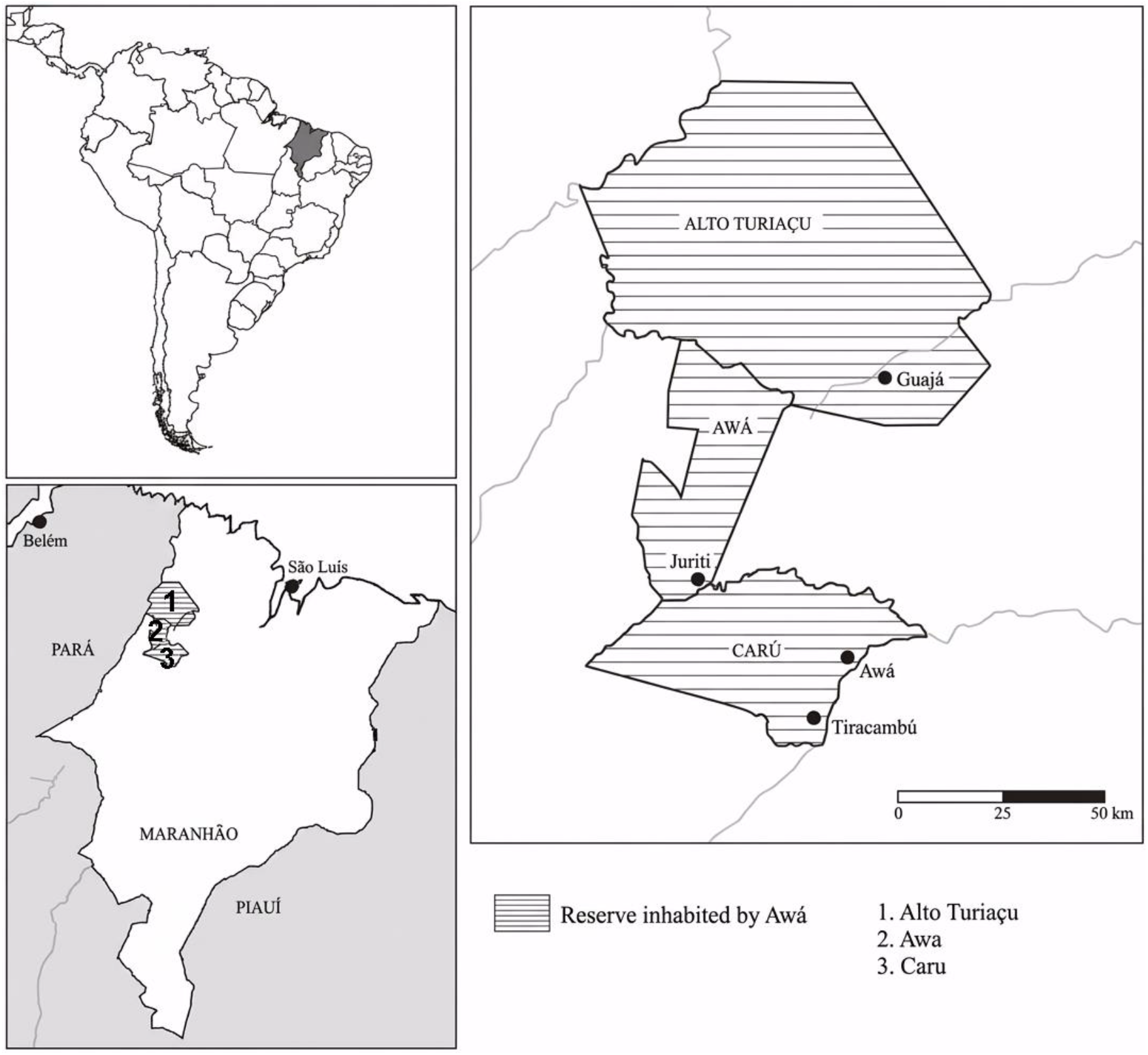
Location of Awá Guajá indigenous reserves in the State of Maranhão. Adapted from Hernando et al., 2011.

This specimen, along with the other materials in the Awá-Guajá collection, is housed and cataloged at the Laboratory of Human Evolutionary Studies (LEEH), affiliated with the Department of Genetics and Evolutionary Biology at the Institute of Biosciences of the University of São Paulo.

Identification was performed through morphological comparisons with specimens housed in scientific collections, including the Renato Kipnis Collection at LEEH and the Mammal Collection of the Zoology Museum of the University of São Paulo (MZUSP). The works of Paula Couto (1979), McBee & Baker (1982), Redford & Wetzel (1985), Feijó et al. (2018), Chahud (2021a), Chahud et al. (2022), and Baldavira Coutinho & Chahud (2026) were also consulted.

## RESULTS

In the Maranhão Amazon region, the family Chlamyphoridae is represented by the species *Priodontes maximus* (Kerr, 1792), *Cabassous unicinctus* (Linnaeus, 1758), and *Euphractus sexcinctus* (Linnaeus, 1758), as reported by Oliveira et al. (2011a, 2011b). The occurrence of *E. sexcinctus* was interpreted as a possible indicator of environmental degradation by Oliveira et al. (2011a, 2011b), who considered the hypothesis that the Awá-Guajá would not hunt this species in anthropized areas or near human settlements. Upon identifying the material as *E. sexcinctus*, Chahud (2022) suggested that the specimen might have originated from hunting activity, as might the other *Dasypus* specimens found in association with it. However, the occurrence of only one specimen of this species led the author to consider other possibilities, such as the occasional presence of an individual near the collection site. Additionally, given the location of the study area near the eastern boundary of the Maranhão Amazon region, the author proposed that the occurrence of *E. sexcinctus* might reflect the influence of vegetation formations associated with the Cerrado in the late 1980s.

This interpretation, however, finds no direct support in the other studies conducted on the Awá-Guajá osteological collection (Chahud, 2019, 2020a, 2020b; Ferreira Figueiredo et al., 2022; Praita et al., 2023a, 2023b). Furthermore, considering the possibility that the specimen may not actually correspond to *E. sexcinctus*, its taxonomic identification should be reassessed. In this context, if the initial species-level identification is incorrect, it is possible that community members did not recognize the animal as belonging to this species, which could explain its inclusion among the animals hunted.

The material originally assigned to *E. sexcinctus* by Chahud (2022) comprises a complete right femur (Figure 2), corresponding to the specimen analyzed in this study (catalog number E27S18/09), as well as a burned femoral fragment bearing cut marks. Chahud (2022) interpreted this second element as potentially corresponding to the left femur of the same individual, based on the similarity in size between the two elements and the absence of other equivalent Chlamyphoridae skeletal elements in the Awá-Guajá collection. However, the fragment’s poor state of preservation compromised the reliability of its taxonomic identification at the species level and limited the acquisition of additional anatomical information. For this reason, the fragment was not directly included in the analyses conducted in the present study and was considered only a possible contralateral element associated with specimen E27S18/09.

**Figure 2.**
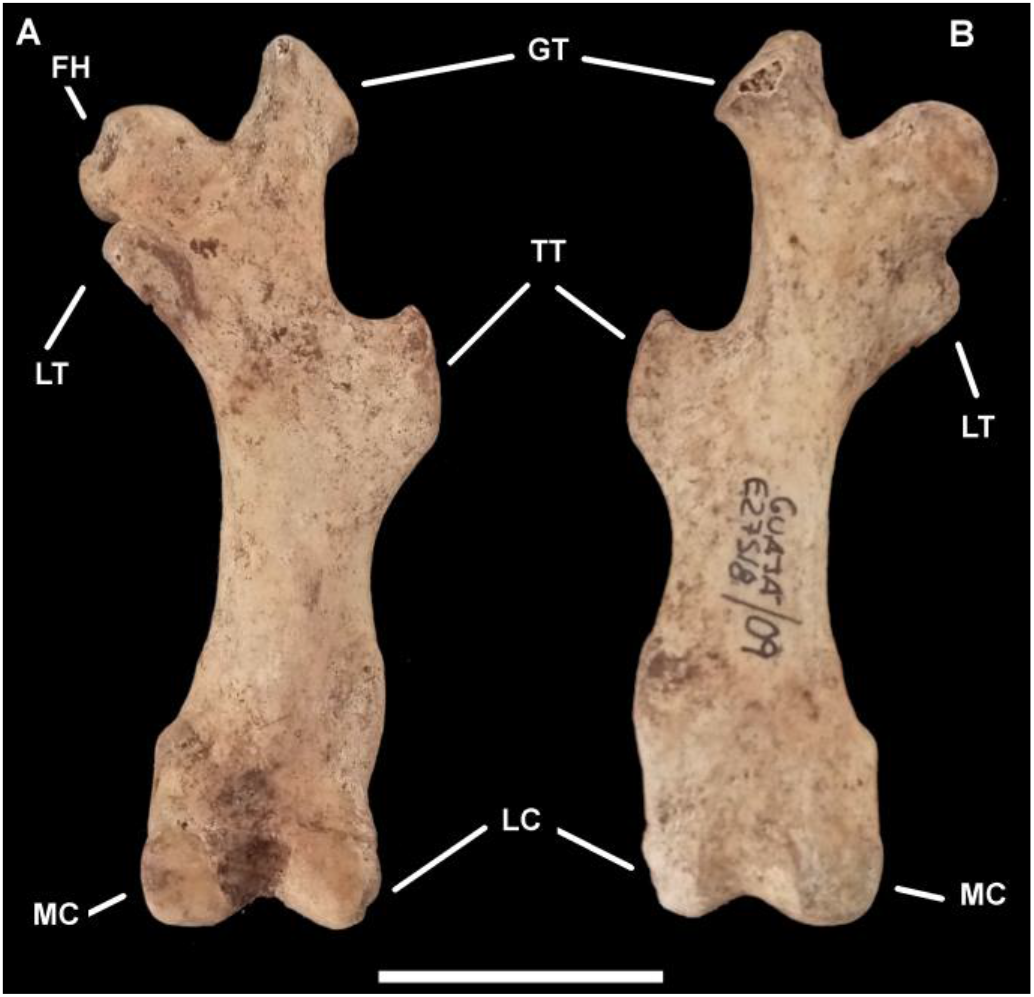
Right femur E27S18/09. A) Posterior view B) Anterior view. Abbreviations: FH - Femoral head; GT - Greater trochanter; LT - Lesser trochanter; TT - Third trochanter; LC - Lateral condyle; MC - Medial condyle. Scale bar: 20mm.

### Remarks

The analyzed femur exhibits ambiguous external morphology, displaying characteristics that allow for the identification of morphological similarities, particularly with specimens belonging to the genera *Chaetophractus, Cabassous*, and *Euphractus*. The fusion of the epiphyses, together with the absence of evidence of pathological alterations, traumatic deformities, or other conditions potentially capable of compromising bone development, such as dwarfism, suggests that the specimen belonged to an adult individual with no bone deformities. Nevertheless, its dimensions indicate dimensions considerably smaller than those of adult representatives of the genera *Cabassous* and *Euphractus* (Table 1).

**Table 1.** Cingulata measurements. Abbreviations: BD: breadth of the distal end; BP: breadth of the proximal end; DC: depth of the caput femoris; DD: depth of the distal end; DP: depth of the proximal end; GL: greatest length (Adapted from Von den Driesch, 1976 and Loponte et al., 2019), *measurements taken without the epiphyses. measurements in mm.

| Specimen | GL | BP | DP | BD | DD | DC |
| --- | --- | --- | --- | --- | --- | --- |
| **E27S18/09** | 62.88 | 22.32 | 9.86 | 17.7 | 11.61 | 7.81 |
| *Euphractus sexcinctus* Adult<br>(CRK no number) | 96.4 | 35.1 | 19.1 | 23.9 | 20.9 | 12.6 |
| *Euphractus sexcinctus* de<br>Lagoa Santa subadult*<br>CVL2- 7000 | 74.39 | 32.12 | 15.27 | 22.71 | 11.8 | -- |
| *Euphractus sexcinctus*<br>Subadult*<br>(CRK no number) | 73.3 | 31.6 | 14.8 | 23.06 | 11.5 | -- |
| *Cabassous tatouay*<br>MZUSP 7665 | 71.7 | 26.0 | -- | 25.6 | -- | -- |
| *Dasypus*<br>Guaja<br>N2E40/02 | 90.15 | 26.06 | 25.00 | 19.67 | 21.38 | 9.34 |
| *Dasypus*<br>Guajá<br>Feat2/68 | 88.33 | -- | -- | 20.76 | 21.3 | 9.5 |

The greatest femoral length is comparable to that recorded for adult individuals of the genus *Tolypeutes* (Figure 3E), but is shorter than that observed in a subadult specimen of *Euphractus* (Figures 3A and 3B), even when the dimensions of the bone without the epiphyses are considered (Figure 3B; Table 1). However, despite this dimensional similarity to *Tolypeutes*, the analyzed specimen exhibits differences in external morphology, particularly in the arrangement and overall configuration of the trochanters, which limits the possibility of a taxonomic association with that genus.

**Figure 3.**
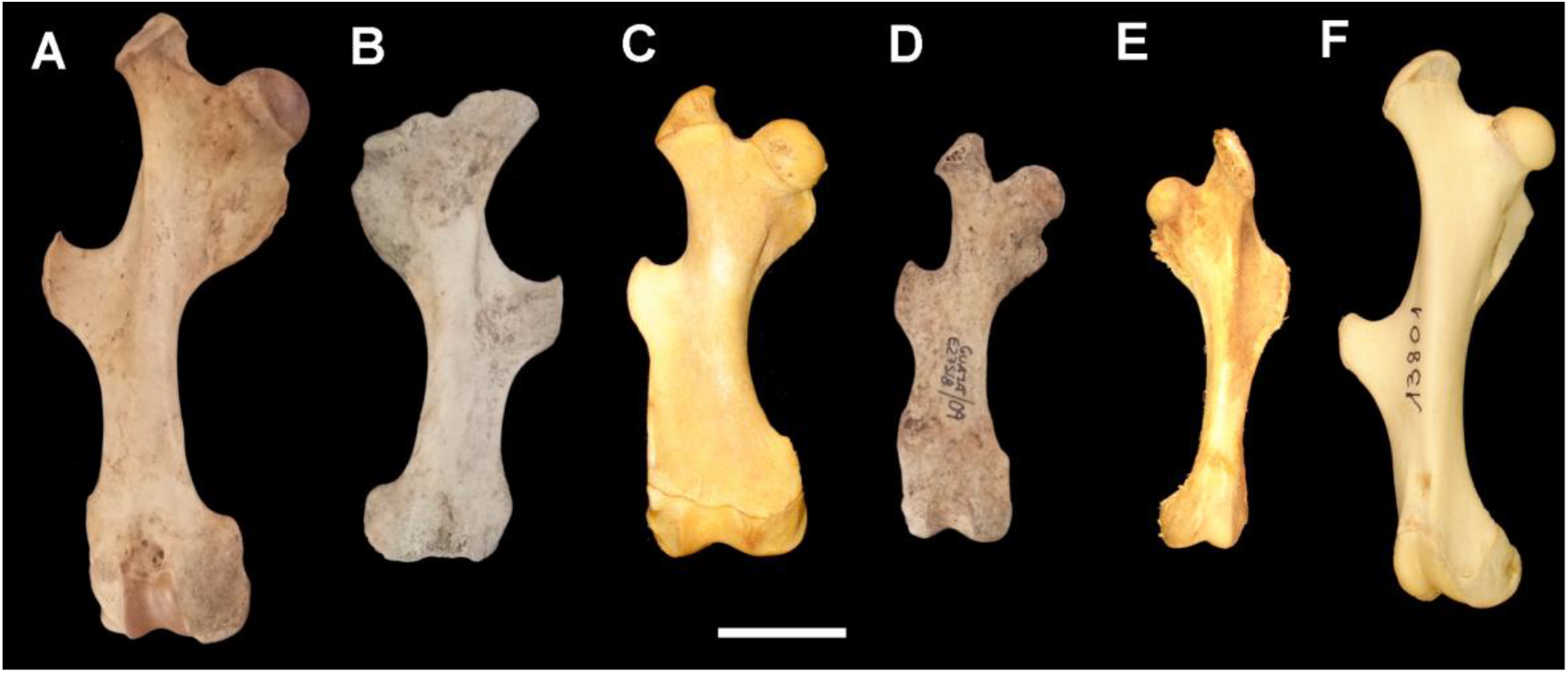
Femurs of Cingulata currently living in Brazilian territory. A) Adult *Euphractus sexcinctus* (Renato Kipnis Collection, no number); B) Subadult *Euphractus sexcinctus* (Renato Kipnis Collection, no number); C) Adult *Cabassous tatouay* Desmarest, 1804 (MZUSP 7665); D) Specimen E27S18/09 from the Awá-Guajá Collection; E) Adult *Tolypeutes tricinctus* Linnaeus, 1758 (MZUSP 3136); F) *Dasypus novemcinctus* Linnaeus, 1758 (MZUSP 13801). Scale bar: 20mm.

The distal end of the femur has a width and external morphological configuration similar to those observed in specimens of *Chaetophractus* and *Euphractus sexcinctus*. In contrast, the overall width of the specimen is considerably smaller than that recorded in representatives of *Chaetophractus* and is also smaller than the width of the distal end observed in *Cabassous* (Figure 3C). On the other hand, the femur has greater transverse dimensions than those found in representatives of the genus *Dasypus*, indicating morphological differences relative to this genus.

The lesser trochanter is relatively elongated and prominent, morphologically resembling the condition described for specimens of *Chaetophractus* from the southern and southwestern regions of South America (Vizcaíno et al., 2003). This shape differs from that observed in representatives of *Cabassous* and *Euphractus*. Additionally, the morphology of this structure differs from the condition observed in *Dasypus* (Figure 3F), showing greater affinity with the morphological pattern described for representatives of the family Chlamyphoridae (Figures 3A–3E).

The distance between the greater trochanter and the third trochanter is closer to the condition observed in *Euphractus* than to that recorded in *Cabassous*. However, the third trochanter has a length that is relatively wider than in *Euphractus*, a characteristic that establishes greater morphological similarity with specimens of the genus *Cabassous* (Figures 3C and 3D). This combination of attributes highlights the occurrence of shared morphological characteristics among different taxa, making identification at the generic level difficult based solely on the available material.

Considering the suite of morphological and metric characteristics evaluated, the specimen can be assigned to the family Chlamyphoridae. However, the observed combination of characters does not allow for an unequivocal assignment to any of the genera compared. Therefore, taxonomic identification should remain restricted to the family level, as assignment to a specific genus would require additional morphological evidence. The recovery of additional osteological material, preferably in the form of a complete skeleton or additional anatomical elements that would allow for a broader morphological comparison, could contribute to a more precise taxonomic identification and to the assessment of the specimen’s affinity with the taxa considered.

## DISCUSSION

Analysis of other species found in the Awá-Guajá Collection provides no evidence that the region was degraded in 1990, and it is likely that forest still predominated in the landscape. Among the animals typical of Amazonian forest environments that have undergone no significant modification, the sloth *Choloepus didactylus* Linnaeus, 1758, stands out, as it is abundant in the collection (Chahud, 2022). Another relevant record is the cervid *Passalites nemorivagus* F. Cuvier, 1817, whose occurrence in sympatry with *Subulo gouazoubira* Fischer, 1814, a cervid typical of the Cerrado and open areas, has been documented at few locations, including this region (Ferreira Figueiredo et al., 2022). In both cases, the Awá-Guajá Indigenous Land represented the eastern limit of the distributions of these species in the state of Maranhão (Chiarello & Plese, 2014; Rossi & Duarte, 2016).

As mentioned earlier, *Euphractus sexcinctus* occurs in the state of Maranhão, including the region inhabited by the Awá-Guajá people (Abba et al., 2025). However, historically, this species has not been a hunting target for the community (Oliveira et al., 2011b), a finding corroborated by the present study. This suggests that the specimen analyzed may not have resembled a typical specimen of *Euphractus sexcinctus*, as the community would also have avoided hunting such an animal.

According to Oliveira et al. (2011b), the only armadillos hunted by the Awá-Guajá community belonged to the genus *Dasypus*. However, it is important to note that these authors’ observations were made approximately 20 years after the Awá-Guajá Collection was collected in 1990, meaning that possible behavioral changes may have occurred during that interval. A possible change in hunting preferences was also observed in relation to wild pigs: in 1990, *Dicotyles tajacu* Linnaeus, 1758, was more strongly represented in the collection (Ferreira Figueiredo et al., 2022), whereas Oliveira et al. (2011b) observed a preference for the larger species, *Tayassu pecari* Fischer von Waldheim, 1814. It is important to note that the *D. tajacu* specimens examined by Ferreira Figueiredo et al. (2022) in the Awá-Guajá Collection corresponded to larger forms found in the Amazon region. A large number of specimens of the sloth genus *Bradypus* Linnaeus, 1758, and of *Tamandua tetradactyla* Linnaeus, 1758, were also recorded in the collection (Chahud, 2022), but these taxa were not recorded in the studies by Oliveira et al. (2011b). In contrast, these authors observed a specimen of *Myrmecophaga tridactyla* Linnaeus, 1758, which was not recorded in the material collected in 1990.

None of the studies considered reported that *Cabassous unicinctus* or *Priodontes maximus* had been hunted by the Awá-Guajá community (Oliveira et al., 2011b; Prado et al., 2012; Chahud, 2022), although both species occur in the region. This suggests that the Awá-Guajá either did not show a preference for hunting these species or that the species were locally rare. The latter hypothesis may apply to *Priodontes* (Oliveira et al., 2011a, 2011b), but not necessarily to *Cabassous*, which is classified as Least Concern (Anacleto et al., 2014) and is considered common in the region. However, *Cabassous* may have been treated similarly to *Euphractus* because of the morphological similarities between the two genera. Considering that similar confusion may also have occurred with the specimen under study, it is not possible to determine whether it represents a distinct variety of Chlamyphoridae, whether its capture resulted from an accidental hunting event by the Awá-Guajá community, or whether it corresponds to a distinct animal whose hunting would not be subject to the same cultural restrictions. It is also not possible to rule out the hypothesis that this form of Chlamyphoridae was actually rare in the region.

The presence of only one femur with distinct morphological characteristics does not provide sufficient evidence to classify it as a new taxon or even as a distinct variety of Cingulata. However, this material suggests the possibility that a form of Chlamyphoridae with distinctive characteristics existed, or may still exist, in the Caru Indigenous Territory, in the state of Maranhão.

This interpretation is justified by cases involving recently proposed species whose taxonomic validity was subsequently questioned or debated. Among the most relevant examples from the Amazon region is the large form of *Dicotyles tajacu* that was assigned a separate species classification as *Pecari maximus* by Roosmalen et al. (2007), but was later disregarded on the basis of genetic studies (Góngora et al., 2007; 2011). Another example is a small tapirid identified as *Tapirus kabomani* by Cozzuol et al. (2013), whose validity has also been questioned for similar reasons (Voss et al., 2014; Ruiz-García et al., 2016).

Unfortunately, it was not possible to perform a genetic analysis of the specimen. However, even if such an analysis had been performed, it might not have been sufficient to identify a new species or even a new variety precisely, given that similar analyses have not been sufficient to establish the taxonomic validity of other proposed taxa. For this reason, the specimen is retained as Chlamyphoridae indeterminate in the present study, pending the identification of more complete specimens that could help clarify its taxonomic position.

## FINAL CONSIDERATIONS

The morphological assessment of femur E27S18/09, housed in the Awá-Guajá Ethnographic Collection, revealed anatomical features consistent with the family Chlamyphoridae. Nevertheless, the characters examined do not provide sufficient evidence to confidently assign the specimen to any of the genera currently recognized within this family. Although some anatomical structures exhibit morphological similarities to those observed in representatives of *Chaetophractus, Cabassous*, and *Euphractus*, the combination of characters identified, considered alongside the femur’s small dimensions, does not support a reliable taxonomic assignment at the generic level. Accordingly, in view of the available evidence, retaining the identification at the family level represents the most defensible taxonomic decision.

The examination of additional specimens, preferably those preserving a broader range of anatomically informative features, could expand the comparative framework and enable a more comprehensive assessment of the specimen’s taxonomic affinities. Complementary analytical approaches, including molecular analyses, may also prove valuable, provided that sufficient suitable biological material is available. Such investigations could refine the taxonomic assessment of the material and improve our understanding of its significance within the diversity of Chlamyphoridae in the Amazonian region.

The identification of zoological material of scientific interest among residues associated with the food consumption practices of Indigenous communities also highlights the potential of these remains for ethnobiological, zoological, and ecological research. Although originally discarded, these materials can provide relevant information about the local fauna, contributing to knowledge of regional faunal composition and to the formulation of hypotheses regarding the relationships between human communities and biodiversity. In this context, the systematic analysis of faunal remains from ethnographic contexts can expand knowledge of regional zoological diversity and support future investigations into species distributions, human–animal interactions, and biodiversity conservation strategies, while respecting the sociocultural contexts of the communities involved.

